# Novel algorithms for PFGE bacterial typing: Number of co-migrated DNA fragments, linking PFGE to WGS results and computer simulations for evaluation of PulseNet international typing protocols

**DOI:** 10.1101/2020.07.05.188623

**Authors:** Ibrahim-Elkhalil M. Adam, Isam Abdokashif, Asia Elrashid, Hiba Bayoumi, Ahmed Musa, Eithar Abdulgyom, Safaa Mamoun, Sittana Alnagar, Wafaa Mohammed, Amna El-khateeb, Musaab Oshi, Faris El-bakri

**Author notes:** Correspondence: Ibrahim-Elkhalil M. Adam.

## Abstract

**Background:** Standard protocols for Pulsed-field gel electrophoresis (PFGE) were adopted and being used in a global scale for surveillance of many bacterial food-borne diseases. Matched PFGE bands are considered regardless of co-migration of different DNA fragments. Molecular epidemiology is turning toward whole genome sequencing (WGS). Although, WGS results can be digested *In-silico*, PFGE and WGS data are being compared separately. We describe a new image analysis algorithm that enables identification of how many DNA fragments co-migrate during PFGE. We built a database that compare described PFGE results to *in-silico* obtained digestion models (from WGS). Reliability of the method was assessed *in-silico* using novel computer simulation approach. From WGS, 1,816 digestion model (DMs) were obtained as recommended by PulseNet international. Simulation codes were designed to predict PFGE profiles when DMs are separated at 5% PFGE resolution in addition to expected co-migration levels.

**Results:** PFGE simulation has shown that about 35% of DNA fragments co-migrate at 5% PFGE resolution. Similar result was obtained when wet-lab PFGE profiles were analyzed using image analysis algorithm mentioned earlier. When image analysis results were compared to DMs, results returned by <u>geltowgs.uofk.edu</u> database revealed reasonable relatedness to DMs. In terms of number of PFGE typable DNA fragments, 45,517 were typable (representing 46.54% out of 97,801). Previously mentioned typable fragments (in terms of typable sizes) comprised 91.24% of the sum of nucleotides of all chromosomes tested (7.24 billion bp). However, significant variations were shown within and between different digestion protocols.

**Conclusions:** Identification of co-migration levels will reveal the third dimension of PFGE profiles. This will provide a better way for evaluating isolate relationships. Linking old PFGE results to WGS by means of simulation demonstrated here will provide a chance to link millions of PFGE epidemiological data accumulated during the last 24 years to the new WGS era. Evaluation of population dynamics of pathogenic bacteria will be deeper through space and time. Selection of restriction enzymes for PFGE typing will have a powerful *in-silico* evaluation tool.

## Introduction

Pulsed-field gel electrophoresis (PFGE) is a form of Agarose gel electrophoresis used to separate large DNA fragments of bacterial chromosomes(1). Such large DNA fragments are obtained using restriction enzymes that has a rare recognition sequence across bacterial chromosome of interest. Consequently, these fragments are generated in few numbers (2,3). Conventional Agarose gel electrophoresis resolve a maximum band size of 40-50 kbp (4). PFGE on the other hand is capable of resolving fragments up to 2 Mbp in size (5). PFGE has shown exceptional reliability for bacterial strain typing (6). Briefly, a PFGE protocol includes the following steps; genomic DNA of bacterial isolates is extracted in a gel plug, digested by the enzyme and resolved in low temperature-melting Agarose gel using counter-clamp homogenous field gel electrophoresis (CHEF). Ethidium bromide is used to visualize resolved fragments(6). Resolution of CHEF was found to range from 10 to 5%. That means at best conditions DNA fragments differ by <5% will resolve within a single PFGE band (a phenomenon known as fragment co-migration) (7,8). A molecular weight marker is run to calculate retention factors (rFs) and DNA band sizes (BS) of bacterial isolates under investigation (9). The optimum number of bands for proper final conclusions is more than ten and less than 30 band (10,11). DNA methylation was found to result in false negative PFGE profiles for the enzyme SmaI (12). Although PFGE refers to other gel electrophoresis methods including field-inversion gel electrophoresis (FIGE) (13) the terms PFGE is being used as a synonymous to CHEF (14). Several protocols were adopted by PulseNet International for typing chromosomes of some pathogenic bacteria species in order to obtain comparable results at national and international levels (15).

A PFGE profile is not always pure DNA fragments. Artifact bands may appear across digestion profiles due to poor washing of extraction plugs. Incomplete digestion also results in false-positive bands, both artifacts are known as ghost bands (8,16). It was also reported that Plasmid DNA may result in false positive bands across PFGE profiles (17). On the other hand, DNA co-migration results in bands with high pixel densities (18). Band intensity profiles are affected by the quality of Ethidium bromide staining and initial cell concentrations which is crucial for both; clearly visible bands across the same lane and less intensity differences between different lanes (10). Comparison of PFGE fingerprints of bacterial isolates is based on the concept of position tolerance; bands that fall within ± 0.015 value of retention factor (rF) are considered a match. In other words; a band exists or none exists within the range of tolerance of another one. That match is considered regardless of co-migration of different DNA fragments (difference in their lengths is too small to be resolved using CHEF) (11).

Computer-assisted analysis of PFGE images helps investigators to numerically express genetic relationships between isolates based on Dice coefficient of variation. Genetic relatedness of different isolates is graphically represented in a hierarchical clustering similarity tree using un-weighted pair group method with arithmetic averages (UPGMA) (19). An exponential correlation was observed between DNA band sizes and their corresponding pixel densities across PFGE lanes [16]. An algorithm was developed by Warner and Onderdonk to consider pixel densities of common bands as comparison parameter for PFGE profiles. Their algorithm was based on standardized trace quantity (STCs) which expresses a kind of qualitative comparison. They reported differences in STCs that indicates DNA co-migration (18), but their algorithm did not answer the question “how many different DNA fragment co-migrated?”.

Whole genome sequencing (WGS) is among the list of PulseNet international for typing methods. WGS includes chromosomal and plasmid DNA in which the entire DNA content of the bacterial isolate is decoded. In this case, sequence alignment provides the ultimate DNA-based typing method. Different approached for sequence alignment include extended multilocus sequence typing (MLST), k-mers and single nucleotide polymorphisms (SNP). Unlike PFGE, WGS provide more valuable details about antibiotic resistance, virulence associated mutations and accurate species and strain identification. PulseNet international is turning toward using WGS instead of PFGE and multi-locus variable number tandem repeats analysis (MLVA)(20).

Typability conclusions based on position tolerance does not take co-migration into account. In addition to the problem of ghost bands. Results of PFGE and WGS are still being compared separately (21). Development of PFGE protocols for different bacterial species is being done by wet-lab PFGE profiling using different restriction enzymes. Although restriction enzyme selection is being done based on WGS data, but a simulation of PFGE separation that enable prediction of the profile is not available. Percentage of PFGE typable size to total chromosome size is not being taken into account in such experiments. The main reasons in our opinion include but not only limited to: firstly, there is no available method that enable quantitative assessment of how many different DNA fragment co-migrate during PFGE and which bands representing single DNA fragment. Secondly, the fact that band sizes calculated from PFGE profiles are estimations based on the molecular weight markers. While, *in-silico* generated fingerprints are exact numeric values. We have developed a method that may enable calculating the number of DNA fragments in each PFGE band.

In this document we tried to answer the following questions:

Q1. How to calculate number of co-migrated DNA fragments across a PFGE profile and what are possible limitations for such a method?

Q2. If co-migration is quantitatively identified, how to compare such results to in-silico obtained digestion profiles (from WGS) considering the fact that PFGE band sizes are estimation based on DNA ladder’s band size vs. rF exponential equation?

Q3. How to predict a PFGE profiles that show co-migration levels from WGS sequences if an *in-silico* digestion profile is obtained?

Q4. If typable PFGE bands are only considered when they fall within the range of the DNA ladder, how much is the percentage from total chromosome size that the sum of typable fragments represents and if optimum number for proper conclusions is from 10 to30, how to evaluate each digestion protocols taking these parameters into account?

## Methodology

### Calculating factor of co-migration

The correlation between band sizes and their corresponding pixel densities was reported to be exponential (16). This finding is the cornerstone of the entire method upon which we made the following assumptions: 1\In theory, a highly significant correlation coefficient (R2>0.99) can be obtained under the following conditions: a. complete digestion by the enzyme is granted. B. Highly pure DNA is extracted within gel plugs. C. all resolved DNA bands represent a single DNA chromosomal fragment (or the same number of fragments). D. High quality Ethedium bromide staining is granted (saturation of DNA content of each band by the dye). Values of band sizes and their corresponding pixel densities can be treated as a two columns matrix in which the first column contains PFGE band sizes and the other column their pixel densities.

Based on the previously mentioned assumptions we suggest that: a\ in case of multiple levels of co-migrated and single-fragment bands occurring across a single PFGE profile, correlation coefficient of exponential equation of band sizes vs. pixel densities will be reduced to a degree proportionate to mentioned levels of co-migrations. b\ in case of analyzing a PFGE profile that have an optimum number of bands (10 to 30 bands), too high (co-migration) and too low (ghosts) values of pixel densities can be removed to have an equation with R^2^ value =>0.98 (only single-fragment DNA bands will remain). The main guideline for identifying such ‘odd’ fragments is to take into account that pixel density should reduce as band size do.

Accordingly if a band size shows a pixel density that is higher than that of the larger fragment, then it represents co-migration and it should be removed. The exception to this rule is the presence of ghost bands. Ghosts may show a significantly low pixel density that my miss lead the entire calculation. c\ by re-creating a polynomial fit and checking values of R^2^, a significantly high R^2^ value can be obtained (0.995 or more.). d\ by denotation of band sizes of the entire profile into the resulting high-R^2^ valued equation, expected pixel densities calculated represents the assumption that all fragment sizes represents a single DNA fragment (a matrix of three columns is obtained at this point). e\ by dividing observed pixel densities (provided by image analysis software) by their expected ones, a proximate number of co-migrated DNA fragments can be obtained. Since integer number is expected, truncation of Observed PD/Expected PD is necessary. This suggested parameter was named factor of co-migration (FCM).

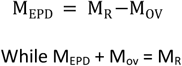

Where M_EPD_; is the two-columns matrix that have R2 value> 0.992 for its correlation equation and from which expected pixel densities will be calculated, M_R_ is the matrix of raw data and M_OV_ is the matrix of odd values of pixel densities.

In order to test the above mentioned method, a meta-analysis of some previously published PFGE results was done (9,16,22). We focused on the DNA marker suggested by Hunter and her team (9) because it run under different conditions across the literature cited (18 different lanes in total). A screen shots for each PFGE image was obtained. Images were saved in .jpg format. Images were imported to GelAnalyzer2010 (23). Gel default colors were set to black DNA bands on white background. Lanes and bands were defined automatically by the software and in some cases, manual modifications were necessary (more frequently for band assignment). Lanes indicated by authors to show XbaI digestion profiles of *S. enterica* serotype Braenderup (strain H9812) where set as DNA ladder for the software. Each band was assigned with its corresponding length in base-pairs. Pixel densities were calculated automatically by the software. Raw data were transferred to Paleontological Statistics (PAST) software package version 4.0. Correlation equation of M_EPD_ matrix was created. Denotations, obtaining values of observed/Expected PDs and truncation to obtain FCMs were all done using Microsoft Office Excel 2007 software package. Mean ± SD and median were calculated from the entire dataset for each band size to get a final FCM-ECSB result using PAST statistics (24). The entire image analysis algorithm described was named factor of co-migration from exponential correlation of single-fragment bands and pixel density (FCM-ECSB). Supplementary material 1 (S1) is a video file showing a complete demonstration for the entire FCM-ECSB method (stream online <u>here</u>).

### In-silico digestion of Whole Chromosome sequences

Chromosomal sequences in this study were chosen from the NCBI Genome database, regardless of randomization or any statistical method for selection. NCBI Genome database was searched for each bacterial species (scientific name + complete genome) that has a standard protocol adopted by PulseNet international. In addition, chromosome sequences for K. pneumonia (which is yet to have a standard protocol) were also included. From table of results returned by NCBI genome database, Plasmids, contiguous and scaffold sequences were excluded. The table of result set was downloaded from the NCBI website in comma separated values file (.csv format). From Replicons column (GeneBank accession numbers) of the result set, Chromosomal sequences were downloaded from the NCBI sequence database in FASTA format. Fragment length for each chromosome were generated considering whether the chromosome is linear or circular using DNADynamo™ software (Blue Tractor Software, North Wales, UK) version 1.0 in default settings. Recommended restriction enzymes for each species were set in separate enzyme boxes (required by the software). Digestion results were exported to text file format (.txt file extension). Raw data were imported to Microsoft office Excel 2007 and refined. Refinement process included removing all metadata for each digestion model except accession numbers. Raw data were transposed into column with accession numbers at the first row. Microsoft Excel files were imported to Microsoft SQL server™ (2014) using SQL server import and export wizard. Data were saved in permanent SQL server database tables. Data are stored in columns representing the length of each DNA fragment for each chromosome sequence in descending order. NCBI GeneBank accession numbers were assigned as a unique identifier (column names) for each digestion model. For data integrity and accurate comparison purposes, table names indicate bacterial species and the restriction enzyme used. Tables of result sets downloaded from NCBI genome database were also imported to the SQL server database to retrieve meta-data from. SQL algorithms were written to compare data uploaded to the system with all models of a specific bacterial species that were generated using the same restriction enzyme and analyzed using the described FCM-ECSB method. Some of the models were used for simulation of PFGE and the assessment of FCM-ECSB method which were shown in this article.

### GelToWGS database algorithms

The database contains 2,420 digestion models. All algorithms were designed to predict number of matched fragments if each DM is run under the same PFGE conditions as test data. To run such simulation, database server is configured to require four parameters from the end user; FCM-ECSB results (Wet-lab PFGE estimated band sizes and corresponding FCMs), which should be uploaded in MS Excel (.xlsx file format), critical co-migration threshold (CCT); it represents resolution quality of PFGE (reported to range from 5 to 10%). Error resulted from running condition (indicated by correlation coefficient of band size vs. rF correlation). Since band sizes estimated from a PFGE profile cannot be treated as exact numeric values, in contrast to in-silico obtained DMs it is almost impossible to get an exact match. Building on this assumption, upper and lower limits for uploaded band size columns are automatically generated. They are filled with data based on a total estimation error (TEE). A final comparison template is required to represent two numbers for each PFGE fragment distanced by a range equals to selected CCT+ marker error. The following equations put theory into a mathematical form:

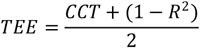

Where TEE; is the total estimation error, CCT; critical co-migration threshold, R^2^; correlation coefficient of marker BS vs. rF correlation, 1; represent 100% R^2^ value. Calculations divided by 2 because to values will be calculated.

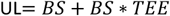

And

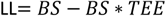

Where UL; is the upper limits. LL; is the lower limits. BS; is a single-column matrix representing wet-lab PFGE band size estimations (uploaded by the end user), TEE; total estimation error.

GelToWGS comparison algorithm will scan each model and count the number of fragments from the model that falls within each range of upper and lower limits across the entire submitted data compared to each digestion model separately. Fragment length is from the genome models, while upper and lower limits are from the image analysis.

When comparing FCMs (from test) to the count (from each model), there are three possibilities: A. Test greater than query; in this case, matched bands = query (the count) B. Test is less than query; matched bands = FCM. C-Equal count and FCM; query count will be considered as matched bands. The computer will calculate the sum of matched bands. Dice coefficient of variation will be calculated to reveal relatedness to each model. Finally, percentage of similarity for both, test and query will be calculated according to the equations:

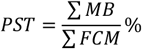

Where PST; is the percentage of similarity to test. *MB;* matched bands. *FCM;* is the factor of co-migration.

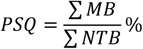

Where PSQ; is the percentage of similarity to query. *MB;* matched bands. *NTB;* is the number of typable fragments (from query) between the biggest and the smallest band size (from PFGE).

The algorithms will also retrieve genome metadata using NCBI accession numbers as a unique identifier that correspond each digestion model. Previously mentioned FCM-ECSB results of *S. enterica* serotype Braenderup strain H9812 (mean values of FCMs truncated to the nearest integer values) where uploaded to the system alongside the following values; 5.2% CCT and 0.002 marker error. Data were compared to 420 whole chromosome sequences of *S. enterica* digested by *Xba*I restriction enzyme (*In-silico* generated PFGE profiles).

### PFGE simulation (WGSToGel)

The target is to evaluate Possibilities for DNA fragments to resolve into single bands and quantitative assessment of co-migration possibilities. Based on the conclusion made by Struelens and his colleagues that DNA fragment co-migration occurs if the percentage of difference between two or more fragments is in the range of 5-10% (6,7). We calculated Dice coefficient % as the difference between a single fragment and the following six fragments after descending order. Mathematically, calculations can be expressed as two parts algorithms:

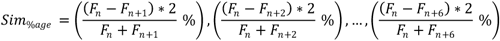

Where Sim_%age_; is a single row matrix (x 6 columns) that represent %age of Dice difference between a single DNA fragment and the following six fragments. F; is the fragments size while n; is the integer number that represents the order of the corresponding fragment size after descending order of the entire digestion profile.

Typable fragments are defined as DNA fragments having sizes within the range of Hunters DNA ladder (20.5 kbp to 1.135 Mbp). Any DNA fragment outside this is automatically excluded from all simulation process. For calculating the Sim_%age_ values for a complete in-silico obtained digestion profile, the same six columns matrix will be created for each typable DNA fragment size. Accordingly, number of rows will equal to typable fragments for each digestion profile, the following algorithm was executed separately for each digestion profile:

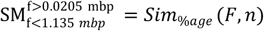

Where SM; is the simulation matrix for fragment lengths of a single digestion profile within the range of 20.5 kpb to 1.135 mega base pair. Sim_%age_; is the equation of single row matrix mentioned earlier. F; fragment size n; is the integer number corresponding the fragment size (F and n are the required parameters for Sim_%age_ equation).

Previously described calculations were made to a total sample size of 1,816 in-silico obtained digestion profiles. Total number of typable fragments is 45,517.

### Band size and FCM simulation

The target was to predict a matrix representing PFGE profile consisting of two columns; band size and corresponding FCM simulations. This algorithm was designed to scan previously described SM matrix and assigning values for band size and FCM based on the following cases:

- **The difference between two fragments > 5%:** the algorithm will give simulated band size the same value of in-silico obtained fragment size. FCM is set to 1 (indicating the expected PFGE band is a single-fragment band). Mathematically, the process can be expressed a follow:

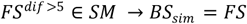 And

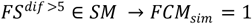

Where FS; is the in-silico obtained fragment size, dif; is Dice percentage of difference (calculated six times), SM; simulation matrix, BS_sim_; expected band size, 1; FCM_sim_ value that indicate no co-migration occurs.
- **The difference between two or more fragments <= 5:** in this case, the algorithm sets simulated band size as the average of co-migrated fragments. While simulated FCM is their corresponding number. The following equations sets mentioned logic into mathematical form:

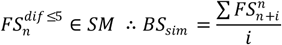 And

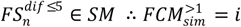

Where FS; is the in-silico obtained fragment size, dif; is Dice percentage of difference (calculated six times), SM; simulation matrix, BS_sim_; expected band size, *n*; order of fragment size, *i*; the number of fragments having <= 5% difference

### Number of typable bands

According to the results of the previously described single and co-migrated bands, simulation algorithm was designed to perform two tasks; firstly, to calculate the total number of bands expected to be visible in actual PFGE assay (single or co-migrated). Secondly, to assign an evaluation rank based on the conclusions described by Van Belkum and colleagues which that the optimum number of bands for proper evaluation of PFGE fingerprints is more than ten and less than 30 (6,11). Accordingly, the same range is ranked as “Optimum” by this algorithm. For models that does not satisfy the cited definition; results were arbitrarily clustered into four different ranks; When number of bands is more than 30; we assigned the label ‘Too much bands’ and when it range from 9 to five, then the rank is “Few bands”. The rank ‘Very few bands’ is assigned when the number of bands is four or three bands. Models that show only one or two bands were labeled as “Non-typable”. This simulation was separately executed for each digestion model.

### Typable size of PFGE to total chromosome size

PFGE typable size is defined as the percentage of the sum of DNA fragments within the range of 1.135 Mbp to 20.5 kbp to the total chromosome size. SQL queries were written to calculate the sum of fragment sizes that satisfy the definition mentioned earlier and calculate the percentage to the whole chromosome size for each model according to the equation:

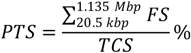

Where: PTS is percentage of typable size. FS; fragment sizes and TCS; is the total chromosome size

## Results

### FCM-ECSB results of Hunters DNA ladder

Mean and median of FCM-ECSB results of Hunter’s marker suggest that out of the 17 PFGE bands; 15 bands represent single DNA fragment. They include bands number 1 to 12, 14 and 16 and 17. Bands number 13 includes two different DNA fragments while band number 15 includes three different co-migrated DNA fragments. Deviations from the mean values occurred in 40 bands (≈13%) out of the whole dataset of 302. When considering standard deviations across band sizes of the marker; the highest SD (±1.2) was shown by band number 15 which we concluded that it represent three different co-migrated fragments. SD for other bands ranged from ±0.0 to ±0.5. We evaluated PFGE profiles based on correlations between band sizes and rF values on one hand and band size and pixel densities on the other; unlike other criteria already adopted in the literature. The results have shown that correlation coefficient of the exponential correlation between BS and rF were highly significant (more than 0.99) and slightly lower between BSs and PDs (between 0.79 and 0.90) in PFGE gels those have 100% FCM matches with medians (same mean values when truncated to integer values); namely Fig. 1 (c) reported by Han and colleagues. The same running conditions were reported to be optimum for typing chromosomes of *k. pneumonia* by the authors. Similar r^2^ values were found in the work reported by Hunter and her colleagues in running conditions recommended for typing chromosomes of both *L. monocytogenes* and *E. coli and Shigella*. Over estimation (as 2 fragments) in one band for each compared to mean values were shown; band #1 and #2 for *L. monocytogenes* and *E. coli-Shigella* protocols respectively. Hunter results for *Salmonella* protocol have shown two deviations in band #8 and #15. Both deviations were over estimations; 2 and 4 for bands #8 and #15 respectively. The exception in Hunter’s work is that bands number 9 and 10 were not separated in gels run under protocols of *E. coli-Shigella* and *Salmonella*. For electrophoresis parameters concluded by Han and colleagues as non optimal; they both showed 32 deviations from the mean (≈10% of deviated FCM estimations). Our estimation criteria mentioned above have shown that R^2^ of BS vs. rF for electrophoresis parameters (EP) a and b were tightly around 0.974 and 0.959 for EP a and b respectively. Which are obviously less when compared to 0.99 shown by previously mentioned protocols. R^2^ values for BS vs. PD for EP a and b showed that in EP a; two lanes showed weak correlations (0.67 and 0.60). It was also obvious that these two lanes included 13 deviations from the mean. Interestingly, they are at the same gel, the other two lanes showed only 4 deviations when R^2^ of BS vs. PD had values around 0.8. For EP b; BS vs. PD correlation was around 0.91. In contrast to EP a; deviations were observed at the end of the marker (smaller band sizes from #12 to #16).

**Fig. (1).**
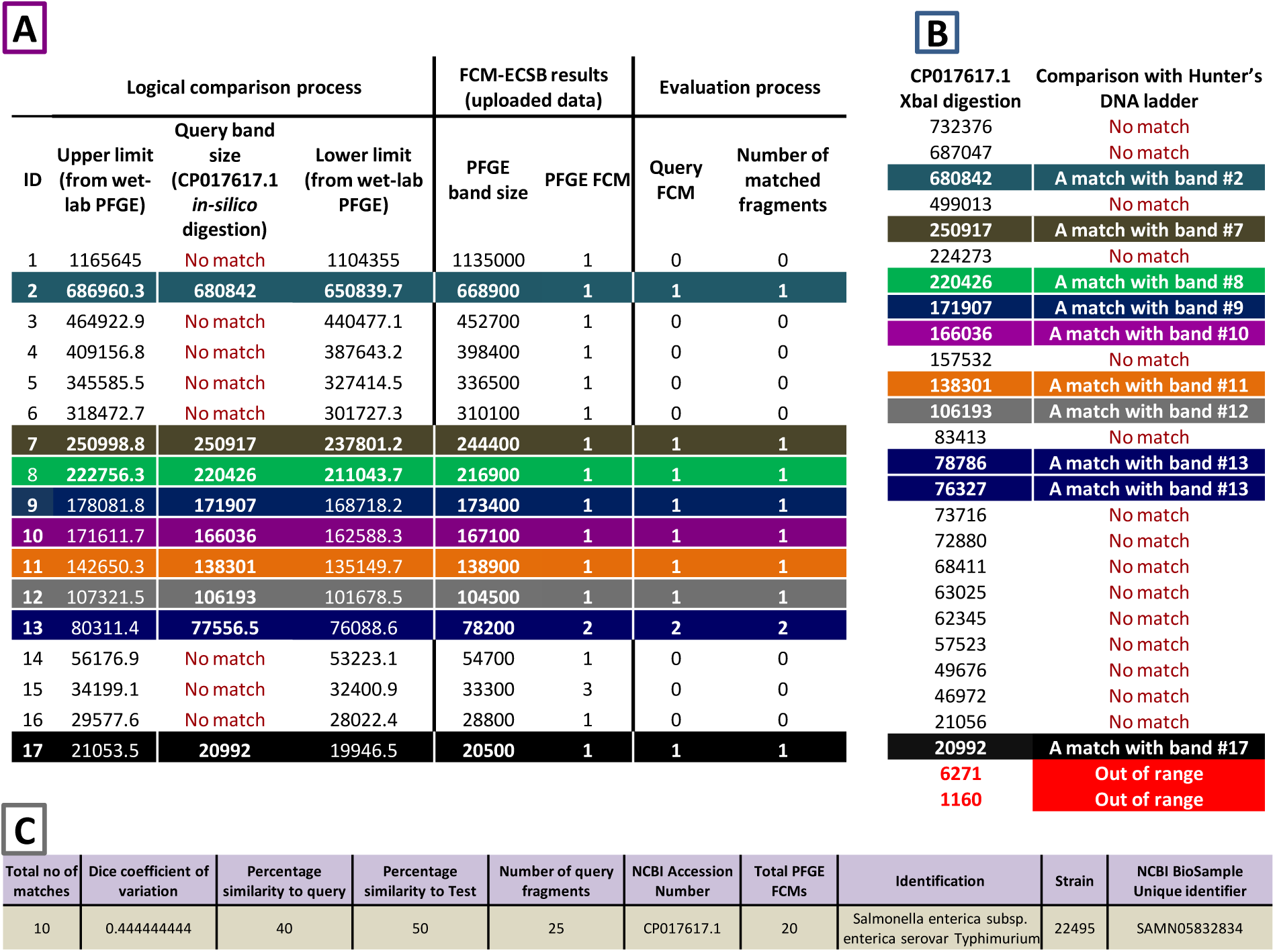
GelToWGS database comparison results: The three tables above show comparisons and evaluation conclusion created by GelToWGS database server when comparing FCM-ECSB image analysis results to digestion profiles of whole chromosome sequences (In-silico driven). PFGE digestion profile of S. enterica serovar Braenderup (strain H9812 digested by XbaI restriction enzyme) was analyzed using FCM-ECSB algorithm in order to reveal number of co-migrated DNA fragments within each PFGE band. FCM-ECSB results (band sizes and corresponding FCMs) where uploaded to the system (analysis parameters: 5% critical co-migration and 0.008 marker error). Uploaded data was compared to in-silico driven XbaI digestion profiles of some S. enterica whole chromosome sequences (310 different digestion profiles in total). *S. enterica* subsp. *enterica* serovar *typhimurium* strain 22495 (GeneBank accession number CP017617.1) has shown the highest number of matched DNA fragments (10). Table A: detailed comparison results showing in-silico driven DNA fragment sizes from CP017617.1 those fall within ranges (upper and lower limits at a distance of about 5%) of actual PFGE band sizes. Matching fragment sizes are indicated by their values and colors (the same as table B). While ‘No match’ indicate that XbaI digested CP017617.1 does not include a fragment size at that range. Columns under the tag ‘Evaluation process’ shows how total matched fragments are considered. Table B: Complete XbaI digestion profile of CP017617.1 showing matched (colored) and non matching (‘No match’)DNA fragment sizes within upper and lower limits of table A. Fragment sizes in red falls outside the typable size of the marker, consequently they are excluded from Dice Coefficient calculations. Table C: shows typability conclusions and metadata of chromosome sequences within NCBI database.

Our FCM-ECSB results showed that separation quality can be evaluated based on correlation between BSs and rFs which largely agreed with conclusions made by the authors. While quality of Ethedium Bromide staining affects detection of co-migration **Table 1**.

**Table 1:**
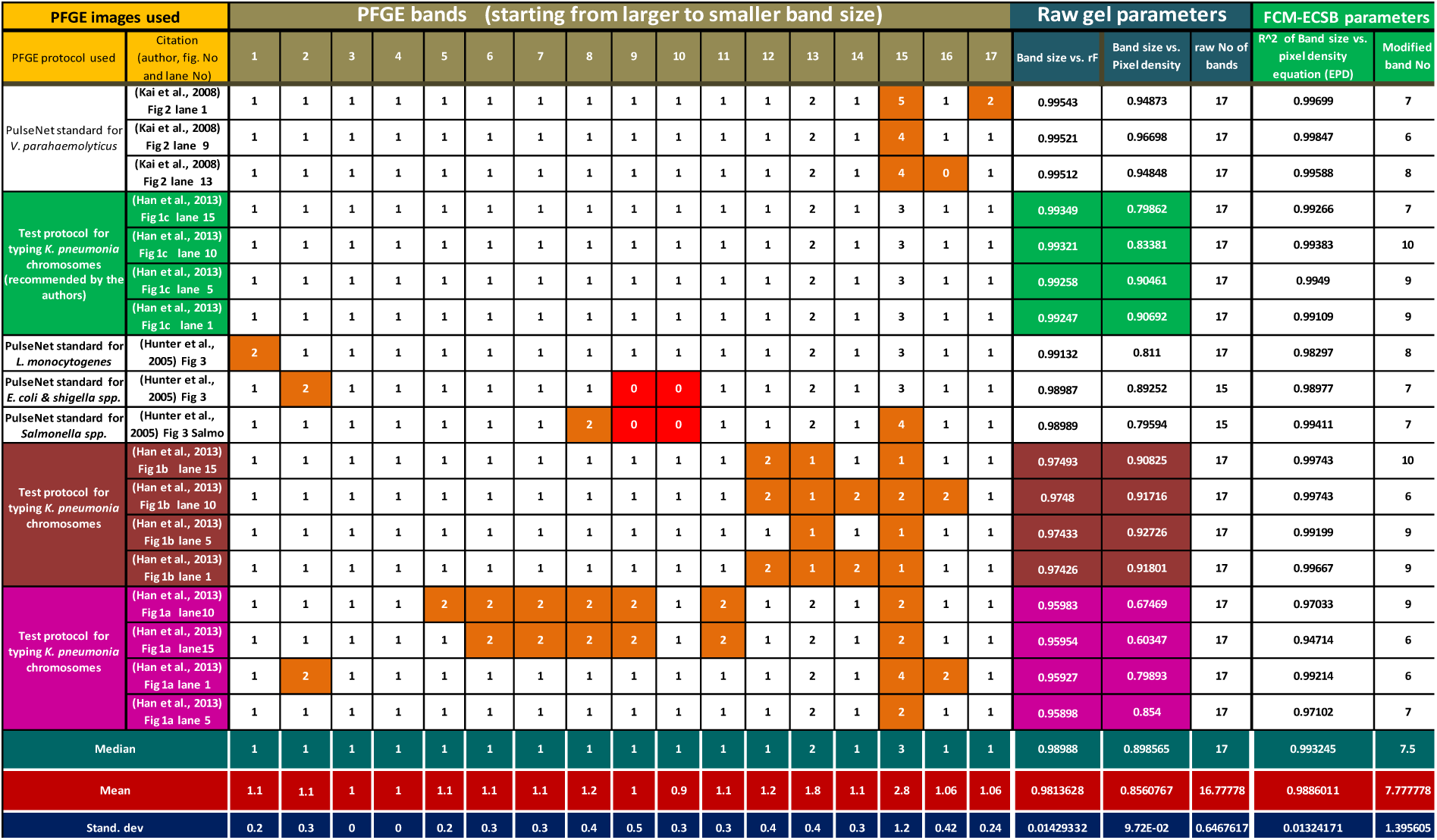
Meta-analysis of *XbaI* digestion of *S. enterica* serotype Braenderup (strain H9812) profiles using FCM-ECSB image analysis algorithm. The table shows number of DNA fragments represented by each PFGE band across *XbaI* digestion profile of *S. enterica* serotype Braenderup (strainH9812)across18differentlanesrununderdifferentelectrophoresisconditions: The first column from left shows citations for each PFGE image (figure numbers) and order of markers lanes from left to right across gel image. Columns under header “PFGE bands” show order of each band of Hunters DNA ladder from the largest band size (1,135 Mbp) to the smallest (0.0205 Mbp. Values highlighted in brown are deviated from truncated values of the average (same as median values). Values highlighted in red were bands not separated (no band size is assigned by the authors). Values under column header “Raw gel parameters” show correlation coefficient of exponential equation of band size vs. retention factor (rF) calculated by GelAnalyzer2010 image analysis software. Column named “Band size vs. pixel density” chows correlation coefficient of polynomial fit created for the entire PFGE profile (column named “raw number of bands” shows total number of marker bands assigned by each author) the correlation between BS and PD was reported to be exponential, but deviations from high R2 is due to co-migration of different DNA fragments. Columns under the header “FCM-ECSB parameters” shows parameters of modified BS vs. PD equation used to calculate expected pixel densities for each band size assuming that selected values represents single-fragment PFGE bands shows how many PFGE bands were chosen to create a polynomial fit between their band sizes and pixel densities to obtain a correlation equation that has a value of R2 shown within column named “R2 values of BS vs. PD equation”. PD values calculated by GelAnalyzer2010 software were divided by expected PDs calculated from FCM-ECSB equation to obtain approximate number of DNA fragments in each band. Since an integer number is expected, values were truncated to the nearest integer value. Results show that 40 band out of 302 (13.24%) deviated from average number of DNA fragments for each band (mean and median values at the bottom of the table).

### GelToWGS comparison algorithm

Out of the 420 digestion model which Hunter’s ladder FCM-ECSB results were compared to, 197 (46.90%) DM showed at least one DNA fragment match with the marker. Total number of typable DNA fragments across the previously mentioned 198 DM was 7,600 typable fragment, out of this figure; matches with Hunter’s DNA marker were 1,461 (19.22%) match. Number of matched fragments ranged from 11 to 4 matched fragments per DM. Taking into account that total number of DNA fragments of the marker is 20, there is no epidemiologically related isolate was shown. *Xba*I digestion profile of *S. enterica* subsp. *enterica* serovar *Typhimurium* (strain 22495) whole chromosome sequence has shown 10 matches and the highest value of Dice coefficient of variation (0.444). Number of typable query fragments of this strain is 25. In contrast to strain SL1344RX of the same serovar which showed 11 matches, that number of query fragments of strain SL1344RX is 51 typable DNA fragment. Percentages of similarities to query for both strains 22495 and SL1344RX are 40% and 21% respectively. **Fig. (1)** Shows details of GelToWGS comparison algorithm upon which previously mentioned conclusions were made. Comparison showed that out of the 17 bands of the marker which represent 20 different DNA fragments, *S. enterica* strain 22495 *Xba*I digestion profile showed a single match with marker’s PFGE band #2, #7 to #12 and #17. Two matches were found with band #13 (Two co-migrated fragments). On the other hand, *S. enterica* strain 22495 *Xba*I digestion profile resulted in 27 DNA fragments out of this figure; two DNA fragments had less than 20,5 kbp (not typable by PFGE). The remaining 25 showed the previously mentioned matches. Supplementary material 2 (S2) shows the details of conclusions mentioned above.

### PFGE simulations

Among the total simulation dataset of 1,816 DM; in terms of size, typable fragments were found to represent 91.24% out of about 7.24 billion bp which represents the sum of nucleotides of whole chromosome sequences tested. In terms of number of chromosomal DNA fragments the total was 97,801. Typable DNA fragments comprised 45,517 fragments (46.54% from total). Our results show that fragment sizes having < 20.5 kbp are actually more than typable ones. Among typable fragments; DNA co-migration when PFGE resolution is 5% reduced number of typable fragments (to expected number of PFGE bands) to 29,430 (35.34% of typable DNA fragments co-migrate). Single-fragment expected PFGE bands comprised 18,225 PFGE bands (61.93% of expected bands). While two co-migrated fragments comprised 7,809 PFGE band which about a quarter of expected PFGE bands (26.53%). Co-migration of three DNA fragments is represented by 2,400 expected PFGE bands (8.15%). Fragment co-migration of 4, 5 or 6 different DNA fragments comprised only 3.39%. Previously mentioned results show that about two thirds of PFGE profiles represent single-fragment PFGE bands. Supplementary material 3 (S3) shows the complete simulation results. **Fig. (2)** Shows a detailed demonstration of how simulation results upon which above conclusions were made. It shows how a PFGE profile is predicted for whole chromosome sequence of *L. monocytogenes* Strain CFSAN023463 (NCBI GeneBank accession number CP012021.1) when it is digested by *Apa*I restriction enzyme.

**Fig. (2):**
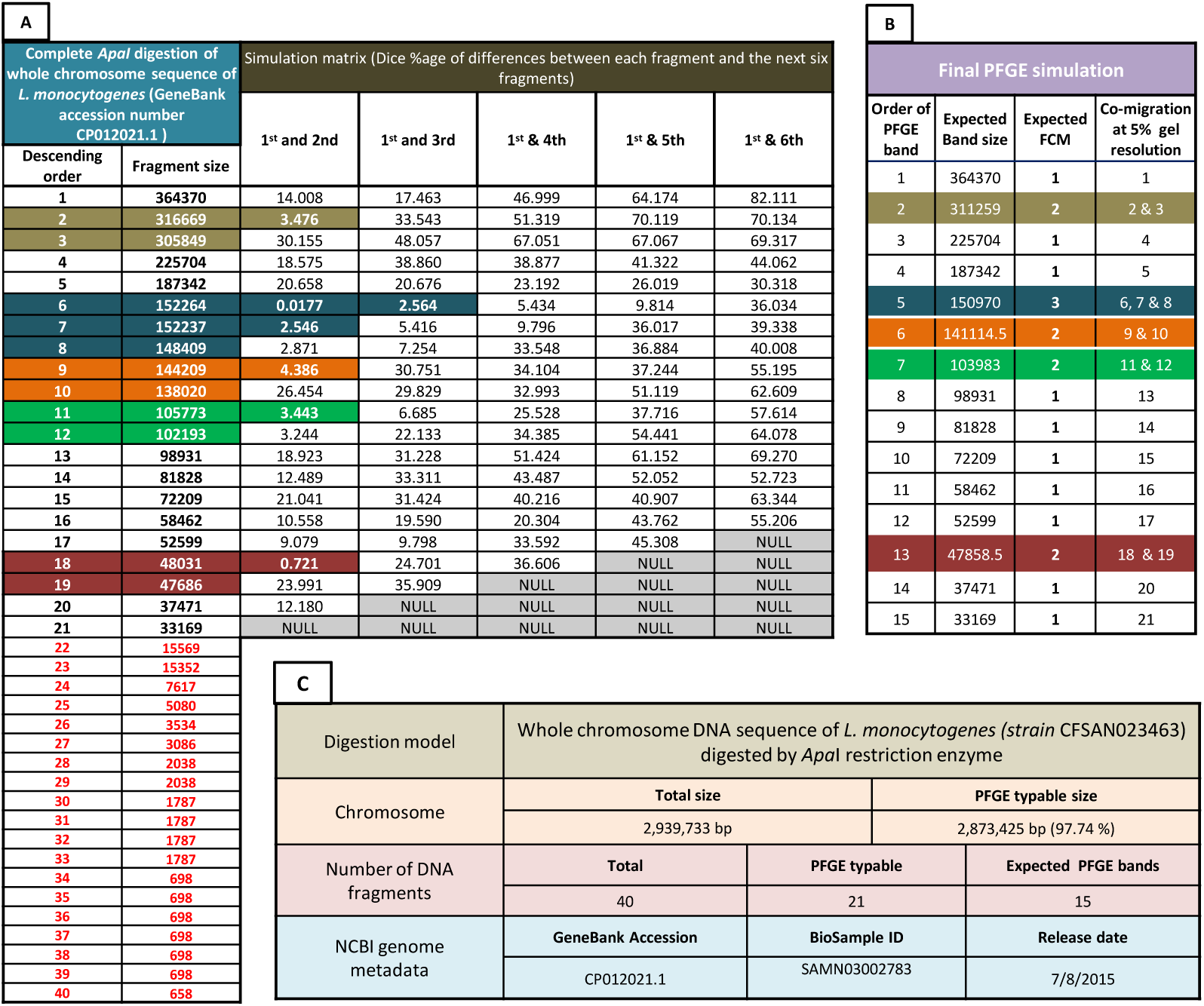
PFGE simulation that show expected PFGE band sizes and corresponding factors of co-migration (FCMs) if the whole chromosome sequence of *L. monocytogenes* (Strain CFSAN023463) is digested by *ApaI* restriction enzyme and run under conditions grant that chromosomal fragments differ by more than 5% are separated: Table A: the first two columns; shows the complete in-silico obtained digestion profile of the mentioned DNA sequence. Fragments shown are in descending order of size regardless of position in the chromosome. Data were obtained using DNADynamo bioinformatics software package. These data simulates digested chromosome sequence in PFGE gel plug before separation. Fragments starting from 22 to 40 are less than 20,5 kbp which is the smallest band size in the widely adopted PFGE DNA ladder reported by S. Hunter and colleagues (values in red). Hence, these fragments were excluded from the simulation. Columns under header text “simulation matrix” show differences between each typable DNA fragment and the following 6 fragments. This difference was calculated as Dice percentage. It is simply calculated by multiplying the absolute subtraction values of each two DNA fragments by 2 and dividing the result by the sum of the two DNA fragments multiplied by 100. this matrix represents possibilities of how likely DNA fragments will co-migrate after PFGE. Factors that determines the final profile is the running distance and quality of running conditions. Table B: shows the final predicted PFGE profile if running conditions and distance were able to separate fragments differ by > 5%. Predictions were made based on simulation matrix (Table A). Colored backgrounds correspond co-migrated DNA fragments across both tables (A and B). Table C: shows over all summaries for digestion profile and metadata of chromosomal sequence. NCBI BioSample ID shows epidemiological metadata.

However, previously mentioned conclusions when each digestion model is considered, significant variation were shown in terms of total co-migration levels. In 18 DMs out of the 26 had single-fragment PFGE bands comprise more than 50%. The rest (8 DMs) had single-fragment bands comprised less than 50%. High percentage of SFBs was shown by *C. jejuni* (*SmaI*) and *L. monocytogenes* (*Asc*I) by 93.71% and 89.24% respectively while, *y. pestis XbaI* and *S. fluxineri* showed SFB by 23.67% and 28.14% respectively. *Y. pestis XbaI* has also shown all co-migration levels tested represented by considerable fractions. The highest percentage of double-fragment bands was shown by *C. botulinum Xba*I and *S. enterica SpeI* by 39.64% and 35.46% respectively. **Fig. (3)** Shows details for each DM.

### Combined PTS and NTP results

Chromosomes with 100% coverage comprised 29 DMs (1.6%). This category showed only optimum (10-30) and few bands (9-5) ranks. Digestion models in this category were dominated by *SfiI* digestions of *V. parahaemolyticus* (14), *AvrII* digestion of *S. enterica* (8). Number of DNA fragments ranged from 5-23. Chromosomes covered by >95%

In addition to *Not*I digestion of *V. cholrae* and *Xba*I digestions of *Y. pestis*. Chromosomes covered by 65-80% (moderately covered) were represented by 179 DM (9.86% from total). 73% of this PTS rank showed few bands (9-5 DNA fragments) and the rest have shown optimum NTB. Interestingly, only three DMs show this PTS rank which resulted from digestion by the restriction enzyme (*Avr*II) for both; *E. coli* and *S. enterica*. Only 4 DMs of *C. botulinum* were also shown this PTS rank. Chromosomes covered by 50-65% (low coverage) comprised only 36 DM out of the entire sample size (1,816). This PTS rank was found to be similar to moderately covered chromosomes in terms of NTB ranks. It also showed Few (86%) and Optimum (14%) NTBs. This PTS rank was also dominated by *Avr*II digestion of *E. coli* and *S. enterica* chromosomes with only two DMs of *L. monocytogenes* (*Asc*I). Poorly covered chromosomes (25-50%) (significantly high PTS) comprised more than three quarters of the entire dataset with 1,385 DM (76.27% from total). this PTS rank showed all NTB ranks from optimum to non typable but optimum NTB rank was the dominant with 1,225 DM (88% of this category) followed by few bands 103 (8%). Models that showed less than 5 DNA fragments comprised only 4%. Interestingly, non typable and Very few bands at this significantly high coverage were shown by 32 and 18 DM respectively. All these models belong to *C. jejuni Sma*I digestion. Small chromosome size of this species is obviously the reason. Chromosomes with high PTS (80-95%) were represented by 126 DM (6.94% of total). They also showed all NTB ranks but unlike significantly highly covered DM, 121 DM showed optimum NTBs (96%). The majority of DMs showed this rank is *Spe*I digestion of *E. coli, S. sonni* and *S. fluxneri*.

are represented by only 17 DMs (the fewest PTS rank). It showed three NTB ranks; optimum, few and very few bands. Un expectedly with this poor PTS, non-typability is not shown. This PTS rank is also dominated by *Avr*II DMs of *E. coli* and *S. enterica*. Chromosomes that show PFGE-typable fragments covering less than the quarter comprised 44 DMs (2.42% of total). As expected, this PTS rank does not include optimum NTBs. In fact, non-typable chromosomes represent 30% of this PTS rank. Very few bands comprised 15 DM (34%) and few bands included 16 DMs (36%). Very poorly covered chromosomes were shown by three DMs. It was dominated by *C. botulimum (Xba*I) followed by *C. jejuni* (*Sma*I) and a single DM of *E. coli* (*Avr*II). **Table (2)** Shows details of association between PTS and NTB ranks.

**Table (2):**
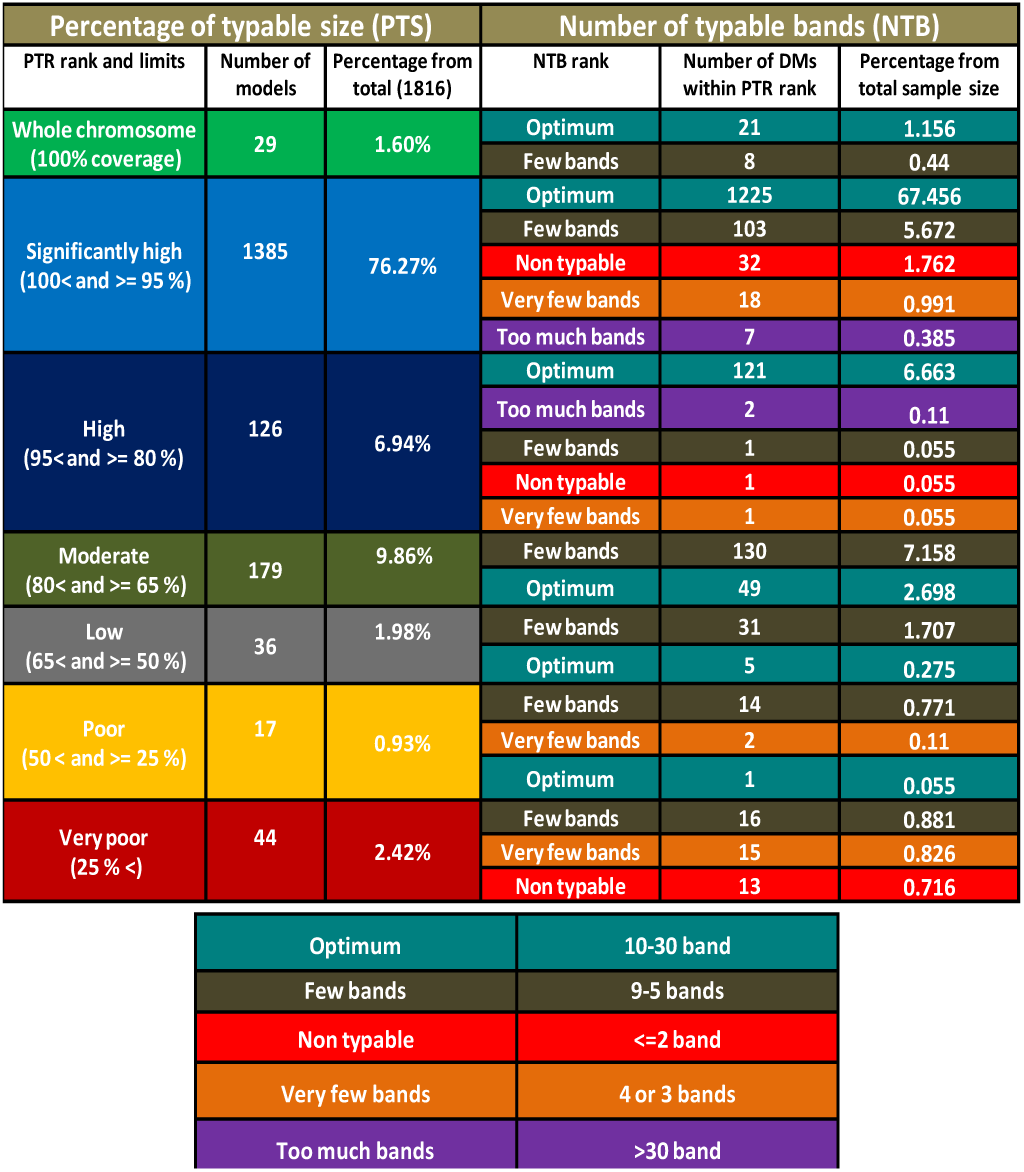
*In-silico* evaluation of currently used PFGE protocols in terms of percentage of typable size to total chromosome sizes (PTSs) and corresponding number of expected PFGE bands for each PTS range. Evaluation and simulation included chromosomal DNA fragments having sizes within the range of 1.135 to 0.205 Mbp and fragments differ by >5% are separated. Among a simulation dataset of 1,816 digestion profile, total number of fragments was 97,801. From this figure; 45,517 (46.54% from total) fragments were typable within the mentioned range. Furthermore, among the typable fragments; DNA fragments co-migration comprised 16,087 (35.34%). Hence, above conclusions were made based on total expected PFGE band comprises 29,430. Generally speaking, adopted PFGE protocols were found to cover major parts of chromosomes tested that; 77.87% have shown PTS coverage at >= 95% including 26 digestion model with 100% coverage. At these significantly high PTSs; number of expected PFGE bands is generally optimum. In contrast to low-PTS digestion models which showed low NTBs.

## Discussion

All described methods in this document do not provide ‘ready to use’ protocols to be adopted. They are simply suggestions those will need lots of experimental evaluation by future research. Here, we would like to summarize some limitations those may have affected our results in addition to some suggested evaluation guidelines for future research.

### FCM-ECSB results

PFGE images used for meta-analysis to demonstrate FCM-ECSB method were obtained in different resolutions. Consequently, pixel densities obtained are greatly affected by resizing of images during production of cited articles. Qualities of cameras fitted with documentation systems used to obtain images are also significant factors. Not to mention quality of Ethedium bromide staining and conversion of images from colored to negative and then black and white. Since manual modifications were made during lane and PFGE bands assignment, human error also might affected our conclusions. That GelAnalyzer 2010 image analysis software calculates pixel densities according to width of bands which were manually modified in many cases. It is also important to mention that separation of bands was incomplete; that intersection of successive bands was observed. The most important drawback in our opinion is the removal of odd pixel density bands, that EPD equation is sensitive especially when number of bands is few. Conclusions made to evaluate PFGE quality depended only on correlation coefficients of both BS vs. rF and BS vs. PD equations.

FCM-ECSB algorithm is obviously sensitive to presence of ghost bands especially those resulted from incomplete digestion. That incomplete digestion affects pixel densities of two bands in addition to presence of a false one.

What about the assumption that selected EPD equation bands represent SFBs taking into account that if they were double or triple fragment bands, they will result in similar high r^2^ values? Two evidences may support our claim; firstly is the simulation results that showed in *Xba*I digestion models of *S. enterica* chromosomes (70 DM in total) SFB comprised 841 expected PFGE bands out of 1551 (54.22%). NTB data mentioned earlier when combined with estimation of typable chromosome size calculated by the sum of band sizes multiplied by corresponding FCM of hunter’s marker result in 4.7 Mbp. This figure is close to average of typable chromosome size of this digestion model (4.53 Mbp ± 381.8 kbp). But when assuming that Hunter’s ladder bands shows at least 2 DNA fragments (FCMs +1), PFGE typable size increased to 9.27 Mbp. This figure is too high even when compared to average of total chromosome size among our data (4.74 Mbp ± 129.5 kbp). When ignoring co-migration (supposing that each PFGE band is SFB), typable size will be 4.56 Mbp. We conclude that FCM-ECSB method may be evaluated based on calculating typable size of chromosome based on the mentioned method.

Simulation results suggest that FCM-ECSB method is hard to adopt for digestion models having few bands or those show little SFBs. For example *Y. pestis (XbaI)* and *S. fluxneri (SpeI)*. They showed SFBs by 23.67% and 28.14% respectively. A possible solution might be using a pixel density calibration ladder that show only SFBs or Hunter’s marker SFBs bands in combination with standardized relative percentage described by Warner and Onderdonk [18] to eliminate variations between different lanes.

Last but not least, we did not perform a comparison between multiple FCM-ECSB results. For performing such comparisons, total estimation errors of both PFGE profiles must be taken into account. Obviously, position tolerance will stay as an important parameter and possibly profile resolution of each individual PFGE profile (expressed as CCT).

### Digestion models derived from WCS

Some FASTA sequences contained ambiguous nucleotides which may have masked some recognition and/or restriction sites of enzymes used. DNA methylation which reported to result in false negative bands might also affect our simulation and GelToWGS comparison results. We suggest that invention of new bioinformatics tools those takes into account nucleotide ambiguity and DNA methylation within recognition sequences of these enzymes may increase accuracy.

### GelToWGS database results

The main idea was to predict Dice similarities assuming that query fragments were run under the same conditions as wet-lab PFGE isolate under investigation. So that, parameters of wet-lab PFGE are the critical factors those determine whole conclusions returned by database server. In other words” GelToWGS database results are only as good as wet-lab PFGE work”. The main factor is CCT value selection. It is obscure because it is hard to determine resolution of PFGE. It depends on running distance alongside other electrophoresis conditions. Another important aspect is plasmid contamination that might result in false positive bands. In this case, conclusions made by the database are skewed. Future upgrades to the system will include plasmid digestion models. Although this upgrade itself may result in more complicated conclusions.

### PFGE simulation results (WGSToGel algorithms)

The target of our simulation was to generate results those are similar to FCM-ECSB image analysis algorithm suggested. The most important question is that does Dice percentage of difference used actually express behavior of DNA fragments during wet-lab PFGE? The answer to this critical question requires wet-lab FCM-ECSB confirmation for simulation results shown in supplementary material 3 (S3) or similar data. According to such findings, WGSToGel simulations can be modified in terms of how difference is calculated. But the main logic will probably remain unchanged.

### Possible contributions to Food-borne disease surveillance

Proposed approaches may collectively improve our understanding of population dynamics and evolution of pathogenic bacteria in many aspects. Since adoption of PFGE for outbreak investigations in 1996, hundreds of millions of PFGE records are available. All these years’ comparisons of PFGE findings were ignoring co-migration of different DNA fragments. Our simulation results suggest that about 35% of PFGE profiles representing co-migrated fragments. But due to absence of a method that reveal number of co-migrated fragments, these genetic variations remained hidden. The described FCM-ECSB image analysis algorithm (if standardized) will provide an important tool to look back and reassess previously made epidemiological conclusions. Based on the same approach, PFGE result might be archived in a database that store band sizes, FCM results and electrophoresis parameters those include correlation coefficients of band size vs. pixel densities and BS vs. rF as numerical data. Such database would not only add a third dimension to PFGE images but it will also require a significantly less storage space.

Since epidemiological surveillance is turning toward whole genome sequencing, linking old PFGE data to WGS results by means of simulation demonstrated here (WGSToGel) will provide a chance to take millions of epidemiological data accumulated during the last 24 years into account while approaching the new WGS era. Such comparisons will only differ from upper and lower limits mentioned earlier in that band sizes are the simulations results, CCT is based on the simulation and marker error is from wet-lab PFGE results. On the other hand, while turning process may take some time, new PFGE data could be simultaneously compared to both old PFGE data and new WGS results.

## Author contributions

Study design, mathematical methods (modeling and algorithms), FCM-ECSB image analysis, SQL database and website programming, SQL simulation algorithms, interpretation of results, statistical analysis and initial manuscript writing by I-E A. In-silico digestion and formulation of some mathematical equations by IA. In-silico digestion of whole chromosome sequences was done by the team that included: I-EA, IA, AE, HB, AM, EA, SM, SA, WM, AE, MO and FE.

## Non standard abbreviations

DM: digestion model
BS: band size
PD: pixel density
SFB: single-fragment PFGE band
FCM: factor of co-migration
CCT: critical co-migration threshold
FCM-ECSB: factor of co-migration based on exponential correlation between single-fragment bands and their pixel densities
PTS: percentage of typable size
NTB: number of typable bands.

## Acknowledgements

The authors are grateful to Dr. Faisal M. Fadelmoula; director of the Centre for Bioinformatics and System Biology (Faculty of Science, University of Khartoum, Sudan) for providing a computer facility for training data entry on using DNADynamo™ software and some data entry. The authors appreciate the efforts of the team who validated the bioinformatic basis, mathematical modeling method, database algorithms and website construction. The team includes; Prof. Omran F. Othman (Dept. of Zoology, Faculty of Science, University of Khartoum) Dr. Abd-Elhameed A. Mansur (Dept. of Computer and information technology, Faculty of Mathematics and computer Science, University of Khartoum), Dr. Muhsin H. Abdalla (Dept. of applied mathematics, Faculty of Mathematics and computer Science, University of Khartoum), Dr. Ahmed M. Elsawe (Africa city of technology, Sudan). We also appreciate the useful discussion with Mr. Mahmood M. Abdulhaleem for his valuable advice for GelToWGS SQL database design.

## Supplementary materials

**Supplementary material 1 S1 (video file)** | Screen-recorded video that demonstrate all calculations of FCM-ECSB image analysis algorithm. The video shows analysis of PFGE profile of *E. coli* O157:H7 strain G5244. Web link is <u>here</u>

**Supplementary material 2 S2 (Excel workbook)** | Results returned by GelToWGS database that show comparisons of FCM-ECSB results of Hunter’s DNA ladder compared to various *XbaI* digested chromosome sequences of *S. enterica* obtained from NCBI GeneBank database.

**Supplementary material 3 S3 (Excel workbook)** | PFGE simulation results obtained using WGSToGel algorithms.

